# Anterior insular CB1 receptor signaling selectively regulates social novelty and anxiety-related behaviors

**DOI:** 10.64898/2026.03.24.713861

**Authors:** Elena Martín-García, Paula Mut-Arbona, Guilherme Horta, Anna Bagó-Mas, Alejandra García-Blanco, Petri Turunen, Michael J. Schmeisser, Inigo Ruiz de Azua, Beat Lutz, Rafael Maldonado

## Abstract

Several neurodevelopmental disorders (NDDs) are characterized by impairments in social behavior and affective dysregulation. Converging evidence implicates the endocannabinoid system in the control of both behaviors. However, the brain region-specific contribution of cannabinoid receptor type 1 (CB1R) signaling to these NDD-relevant phenotypes remains unclear. The anterior insular cortex (aINS) is a key integrative hub involved in socio-emotional processing and social novelty recognition. Whether CB1Rs within this region are sufficient to regulate behavioral domains disrupted in NDDs remains unclear. Here, we employed a Cre-dependent viral strategy to selectively restore CB1R mRNA expression in the aINS of global CB1R-deficient mice. Region-specific rescue of CB1R in the aINS normalized social novelty discrimination and reduced anxiety-like behavior as compared to mice lacking CB1R, while leaving basal sociability and locomotor activity unaffected. In addition, insular CB1R re-expression modulated repetitive-like behaviors without broadly altering other behavioral domains. These effects were observed in the absence of off-target expression, supporting the specificity of the genetic manipulation. Our findings demonstrate that CB1R mRNA expression within the aINS is sufficient to regulate distinct socio-emotional and repetitive behavioral domains. These results identify the aINS as a critical CB1-dependent modulatory node and provide mechanistic insight into how region-specific endocannabinoid signaling contributes to behavioral phenotypes relevant to NDDs.

## 1. Introduction

Neurodevelopmental disorders (NDDs) comprise a heterogeneous group of conditions arising from disruptions in brain maturation that persist throughout life. These disorders, including attention-deficit/hyperactivity disorder (ADHD), autism spectrum disorder (ASD), intellectual disability (ID), and specific learning disorders, are characterized by impairments in cognition, behavior, and social functioning. Their origins reflect complex genetic–environmental interactions that disrupt the structural and functional organization of the nervous system, affecting domains such as memory, language, motor skills, and social behavior^1,2^. Importantly, individuals with NDDs are at increased risk for psychiatric disorders later in life^3^.

NDDs frequently co-occur and display overlapping phenotypes, particularly deficits in social cognition and emotional processing^4,5^. Such convergence suggests dysregulation of shared neuromodulatory systems. Among these, the endocannabinoid system (ECS) has emerged as a key regulator of neurodevelopment, synaptic plasticity, and circuit homeostasis^6,7^. Altered cannabinoid signaling has been implicated in schizophrenia^11^, Fragile X syndrome^8^, and ASD-related phenotypes^9^ among others, highlighting the ECS as a potential convergent pathway in neurodevelopmental vulnerability.

Endocannabinoid signaling is primarily mediated by cannabinoid receptor type 1 (CB1R), the most abundant G protein–coupled receptor in the brain^10^. CB1Rs regulate excitatory and inhibitory neurotransmission at synapses, positioning them to fine-tune neural circuit dynamics underlying cognitive and social behaviors^11,12^. Despite this widespread role, the region-specific contribution of CB1R signaling to NDD-related behaviors remains poorly defined.

The anterior insular cortex (aINS) is increasingly recognized as a central hub for integrating socioemotional processing, interoceptive awareness, and social recognition memory^13-17^. Anatomically and functionally distinct from the posterior insula, the aINS integrates sensory information with emotional salience through extensive connectivity with limbic and prefrontal regions^13,15,17^. Recent studies demonstrate that local insular circuitry dynamically encodes social novelty and preference^14,18,19^, and disruption of these circuits impairs social recognition. Moreover, converging evidence indicates functional interactions between the ECS and insular circuits in regulating social and affective behaviors^20^.

Despite this emerging evidence, it remains unclear whether CB1R signaling within the aINS is necessary and/or sufficient to regulate behavioral domains commonly disrupted in NDDs. To address this gap, we used stereotaxic viral strategies in genetically engineered mouse models to selectively manipulate CB1R expression in the aINS. Given the transdiagnostic nature of NDD symptoms^21^, we assessed multiple behavioral domains, including social behavior, repetitive behaviors, anxiety-like behavior, depressive-like behavior, and locomotor activity. By evaluating full rescue, global deletion and region-specific rescue of CB1R mRNA expression in the aINS, we aimed to determine the causal contribution of local CB1R expression to behavioral phenotypes relevant across NDDs.

## 2. Material and Methods

### 2.1. Animals

Male mice were housed under standard laboratory conditions in a temperature-controlled (22 ± 1 °C) and humidity-controlled (50 ± 10%) environment under a 12:12 h inverted light/dark cycle (lights on at 19:00). Food and water were available *ad libitum*. All experiments were performed in adult male mice (8 weeks of age) maintained on the same genetic background. The CB1 knockout (STOP-CB1) line carries a loxP-flanked transcriptional STOP cassette inserted into the 5′ untranslated region of the *Cnr1* gene upstream of the ATG start codon, resulting in complete CB1 receptor deficiency, as previously described^22^. To obtain control mice with global CB1 re-expression, STOP-CB1 animals were crossed with EIIa-Cre deleter mice, leading to germline excision of the STOP cassette and generation of a CB1 rescue (CB1-RS) allele that restores CB1 expression in all cells; EIIa-Cre was then bred out, and these CB1-RS, Cre-negative Ella-WT animals are referred to here as control WT. All lines were backcrossed for at least seven generations onto a C57BL/6J background.

### 2.2. Experimental design

To investigate whether CB1 receptor signaling in the aINS is sufficient to regulate behavioral phenotypes, we used a stereotaxic viral strategy in genetically engineered mice. Three experimental groups were established: (1) WT mice injected with AAV-GFP into the aINS, (2) STOP-CB1 mice injected with AAV-GFP (lacking CB1 expression), and (3) aINS-CB1-RS mice, which are STOP-CB1 mice injected with AAV-syn-Cre-GFP to rescue (RS) CB1R expression locally in the aINS (Fig. 1A). Bilateral viral injections were performed stereotaxically at 8 weeks of age, and animals were allowed four weeks for stable viral expression before behavioral testing. Behavioral phenotyping was conducted using a structured battery of tests assessing repetitive-like behavior (marble burying), social behavior (three-chamber social interaction and social memory), locomotor activity (automated locomotor chambers), anxiety-like behavior (elevated plus maze), and depressive-like behavior (forced swim test). Testing was performed in a fixed order progressing from low-to high-stress paradigms to minimize carry-over effects (Fig. 1B). Each test was conducted on a separate day with at least 24 hours between sessions. All experiments were carried out during the animals’ active (dark) phase under red light conditions, and the experimenter was blinded to genotype and treatment group throughout data acquisition and analysis.

**Figure 1.**
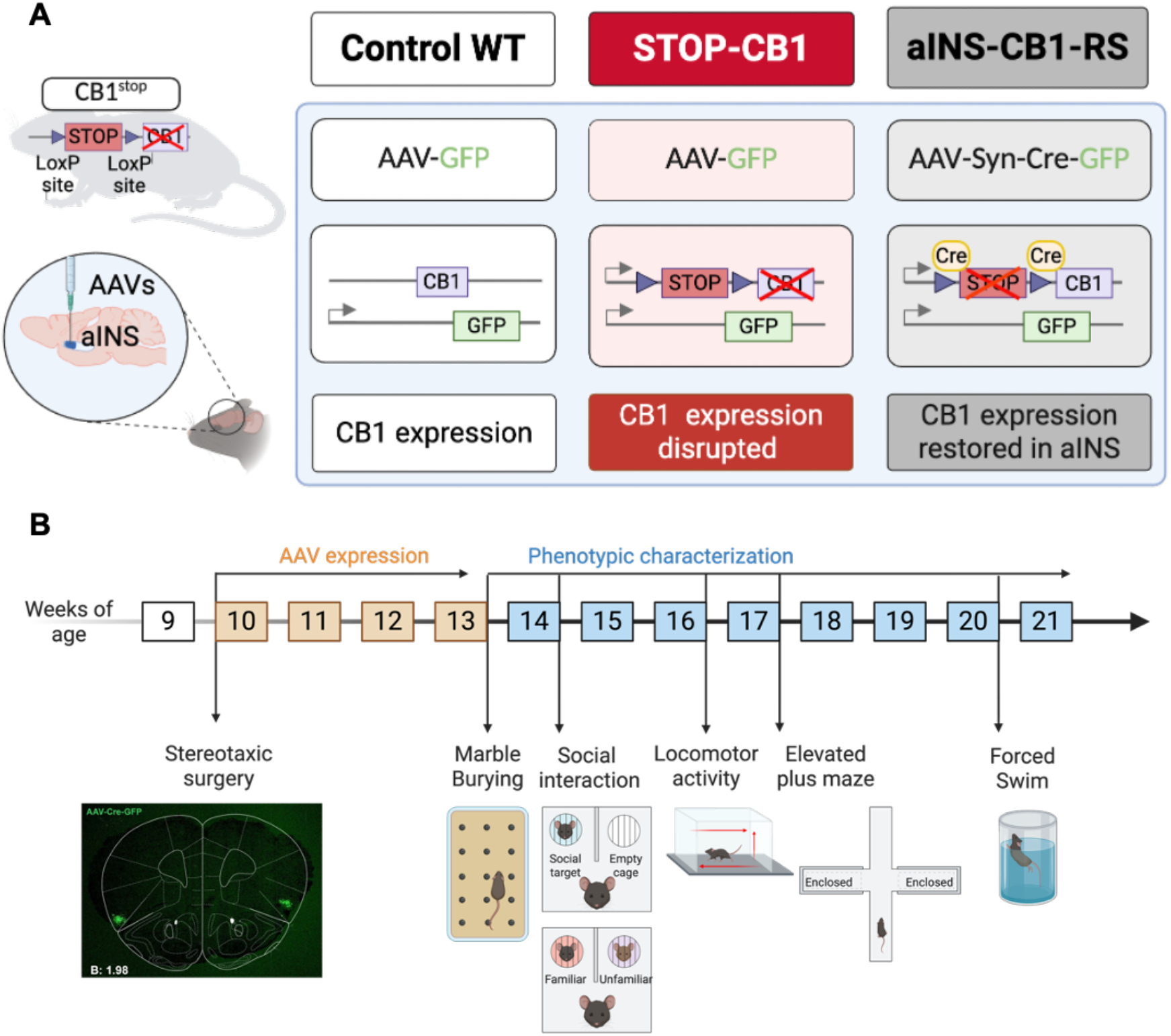
Experimental design and region-specific rescue strategy of CB1 receptors in the anterior insula. **(A)** Schematic representation of the experimental groups and viral strategy. WT mice exhibit normal CB1R expression. STOP-CB1 mice carry a loxP-flanked transcriptional stop cassette inserted upstream of the *Cnr1* coding sequence, resulting in global absence of CB1R expression. In the aINS-CB1-RS group, bilateral injection of AAV-Syn-Cre-GFP into the aINS induces Cre-mediated excision of the stop cassette, restoring CB1R expression selectively within the aINS. **(B)** Timeline of the experimental procedure. Mice underwent stereotaxic surgery followed by a four-week period to allow viral expression. Behavioral phenotyping was subsequently performed in the following order: marble burying, three-chamber social interaction (sociability and social novelty), locomotor activity, elevated plus maze, and forced swim test.

### 2.3. Stereotaxic surgery and viral vector microinjection

To achieve region-specific restoration of CB1 receptor (CB1R) expression in the aINS, we used STOP-CB1 mice carrying a loxP-flanked transcriptional stop cassette inserted upstream of the CB1R coding sequence. In these mice, CB1R expression is globally suppressed until Cre-mediated excision of the stop cassette restores receptor expression locally. For the aINS-CB1-Rescue (RS) group, STOP-CB1 mice received bilateral stereotaxic injections of an AAV vector encoding Cre recombinase under the synapsin promoter (AAV-Syn-Cre-GFP) into the aINS. WT and CB1-KO control animals received bilateral injections of AAV-Syn-GFP into the same coordinates. Surgeries were performed under general anesthesia induced by intraperitoneal injection of ketamine hydrochloride (75 mg/kg) combined with medetomidine hydrochloride (1 mg/kg), both diluted in sterile 0.9% saline. Mice were secured in a stereotaxic frame (David Kopf Instruments, USA), and bilateral injections were delivered into the aINS using 33-gauge internal cannulas connected via polyethylene tubing (PE-20) to a 10 μl Hamilton microsyringe. A total volume of 0.2 μl per hemisphere was infused at a rate of 0.05 μl/min using a microinfusion pump. Following injection, the cannula remained in place for 10 minutes to allow diffusion and minimize reflux before slow withdrawal. Coordinates targeting the aINS were adapted from Paxinos and Franklin’s mouse brain atlas: anteroposterior (AP) +1.9 mm, mediolateral (ML) ±3.5 mm, and dorsoventral (DV) −2.0 mm relative to bregma. Postoperative care included subcutaneous administration of meloxicam (2 mg/kg) for analgesia and gentamicin (1 mg/kg) to prevent infection.

### 2.4. Behavioral Testing

All behavioral procedures were conducted in accordance with institutional animal care guidelines and approved by the Animal Ethics Committee of the Parc de Recerca Biomèdica de Barcelona, CEEA-PRBB and the Generalitat de Catalunya (1649-(PG)EMG-22-0071-P2) and under European Directive 2010/63/EU. See Fig. 1B for the scheduled order of behavioral tests. Behavioral tests were performed with adequate inter-test intervals, and the sequence of testing was optimized to minimize potential order effects.

#### 2.4.1. Marble Burying Test

Repetitive-like behavior was assessed using the marble burying test^23^. Mice were placed individually in transparent plastic cages (26.5 × 20 × 14 cm) containing 5 cm of sawdust bedding with 20 evenly spaced glass marbles (14 mm diameter). After 30 minutes, mice were returned to their home cages, and the number of marbles buried (defined as ≥ two-thirds covered with bedding) was counted.

#### 2.4.2. Three-Chamber Social Interaction Test

Sociability and social novelty recognition were evaluated using a three-chamber social test adapted^24^. The apparatus consisted of three interconnected chambers (20 × 40 × 22 cm each) with clear Plexiglas walls and openings (35 × 35 mm) allowing free movement between compartments. During habituation, test mice were confined to the central chamber for 5 minutes. In the sociability phase of social interaction, a novel conspecific (Stranger 1; C57BL/6NCrl male) was placed inside a wire cage (60 × 60 × 100 mm) in one side chamber, while an identical empty cage was placed in the opposite chamber. Mice were allowed to explore all chambers for 10 minutes. Time spent in each chamber and the number of entries (all four paws inside) were recorded and manually scored. In the social novelty phase, we measured social memory, and a second unfamiliar mouse (Stranger 2) was placed in the previously empty cage for an additional 10-minute session. Preference for social novelty was assessed by comparing the time spent exploring the novel versus the familiar conspecific.

#### 2.4.3. Locomotor Activity

Spontaneous locomotor activity was assessed in individual activity chambers (9 × 20 × 11 cm; Imetronic, France) under low-light conditions (20–25 lux)^25^. Horizontal and vertical movements were automatically recorded over a 120-minute period.

#### 2.4.4. Elevated Plus Maze (EPM)

Anxiety-like behavior was evaluated using the elevated plus maze. The apparatus consisted of two open arms (30 × 5 cm) and two closed arms (30 × 5 × 15 cm) connected by a central platform (5 × 5 cm) elevated 50 cm above the floor^25^. Mice were placed on the central platform facing an open arm and allowed to explore freely for 5 minutes. Behavior was recorded and manually analyzed. The apparatus was cleaned between animals, and illumination was maintained at 10 lux in closed arms and 200 lux in open arms. Time spent in open arms was used as an index of anxiety-like behavior. Total entries in both open and closed arms were recorded.

#### 2.4.5. Forced Swim Test (FST)

Depressive-like behavior was evaluated using the forced swim test^26^. Mice were habituated to the testing room for 30 minutes prior to the experiment. Animals were placed in a transparent Plexiglas cylinder filled with water maintained at 25 °C for 6 minutes. Immobility, defined as floating with only minimal movements necessary to keep the head above water, was quantified during the final 4 minutes. Following testing, mice were dried and monitored in a clean cage placed on a heating pad. Behavior was scored manually.

### 2.5 Immunofluorescence and Imaging

#### 2.5.1. Tissue preparation

Mice were deeply anesthetized and transcardially perfused with 4% paraformaldehyde (PFA) in phosphate buffer (0.1 M, pH 7.5). Brains were post-fixed overnight in 4% PFA and cryoprotected in 30% sucrose at 4 °C. Coronal sections (30 μm) were obtained using a freezing microtome and stored in cryoprotectant solution at 4 °C until processing.

#### 2.5.2. Immunofluorescence

Free-floating sections were rinsed in phosphate buffer and blocked in a solution containing normal goat serum and Triton X-100. Sections were incubated overnight at 4 °C with primary antibodies against Cre recombinase, CB1 receptor, or GFP as appropriate for experimental validation. After washing, sections were incubated with species-specific Alexa Fluor–conjugated secondary antibodies for 2 h at room temperature. Sections were then mounted on gelatin-coated slides using Fluoromount mounting medium.

#### 2.5.3. Confocal imaging and analysis

Confocal imaging was performed using BC43 spinning disk confocal system (Andor Oxford Instruments, United Kingdom). Laser lines of 488 (10% power) and 561 nm (5% power) were used for the fluorescence excitation and the fluorescence emission was acquired using bandpass filters 529/24 and 595/31 for GFP and Alexa Fluor 594-CB1, respectively. The imaging was performed sequentially to minimize the spectral crosstalk. The aINS for both sides of the brain were imaged in using a CFI Plan ApoChromat Lambda D VC ×20/0.8 NA air (Nikon) objective and 4.2 MP Zyla Camera (6.5 µm pixel, 16-bit, 100 ms exposure time for both channels) with a pixel binning size of 2 (effective image pixel size of 609 nm) as 10 μm thick z-stack with a slice interval of 0.4 μm. In total, four brain sections were imaged per mice for three WT, two STOP-CB1 and four aINS-CB1-RS conditions.

Quantification of CB1-AF594 filamentous structures was performed by analyzing the fractional filament/tissue area for CB1-AF594 fluorescence channel. In the first step, the z-stacks were deconvolved using the image acquisition software Fusion (1.1.0.1), followed by 3D gaussian filtering (σ = 1px) and performing maximum intensity projection (MIP). Segmentation of the filamentous and tissue area was performed using Ilastik random-forest machine learning based pixel classification^27^. Training image set consisted of five images representing all three experimental conditions (WT/STOP-CB1/aINS-CB1-RS). For training, all “edge” and “texture” based feature filters (σ = 0.7-10) were used. Pixel classifier was trained by manual labelling of “filaments”, “tissue” and “outside tissue” areas of from all the images of the training image set until the classifier performed the segmentation visually in satisfactory quality. Fifty percent probability from the probability map was used as segmentation threshold.

The further processing was performed using custom made macro in Fiji imageJ^28^. The segmentation mask for “filaments” was processed by filtering out small and circular objects (size < 15 px, circularity >0.5). The “Tissue” mask was processed with median filter (σ = 2px) and filter out small objects (size < 100 px). The final readout was the fraction of “filament” and “tissue” area over the imaged area. All analyses were conducted blinded to genotype and treatment group.

### 2.6. Statistical Analysis

Statistical analyses were performed using GraphPad Prism (version 11). Data are presented as mean ± standard error of the mean (SEM), with the number of animals (n) indicated in the figure legends and figures. Normality was assessed using the Shapiro– Wilk or Kolmogorov–Smirnov test, as appropriate. For comparisons between two groups, parametric data were analyzed using unpaired two-tailed Student’s t-tests, whereas these non-parametric comparisons were analyzed using the Mann–Whitney U test. Comparisons involving more than two groups were analyzed using one-way ANOVA followed by Bonferroni post hoc correction for parametric data. Two-way repeated-measures ANOVA followed by Bonferroni or Sidak’s post hoc tests was used when appropriate. Statistical significance was defined as *p < 0.05, **p < 0.01, ***p < 0.001; n.s., not significant. All graphs display mean ± SEM.

## 3. Results

### 3.1. Region-specific restoration of CB1R expression in the anterior insula

To determine whether CB1R expression in the aINS is sufficient to regulate behavioral phenotypes, we first validated the anatomical specificity and effectiveness of our viral rescue strategy (Fig. 2). STOP-CB1 mice carrying a loxP-flanked transcriptional stop cassette received bilateral AAV-Syn-Cre-GFP injections into the aINS to generate the aINS-CB1-Rescue group, whereas WT and STOP-CB1 control animals received AAV-GFP. Coronal sections confirmed accurate targeting of the aINS according to the Paxinos and Franklin atlas (Fig. 2A–G).

**Figure 2.**
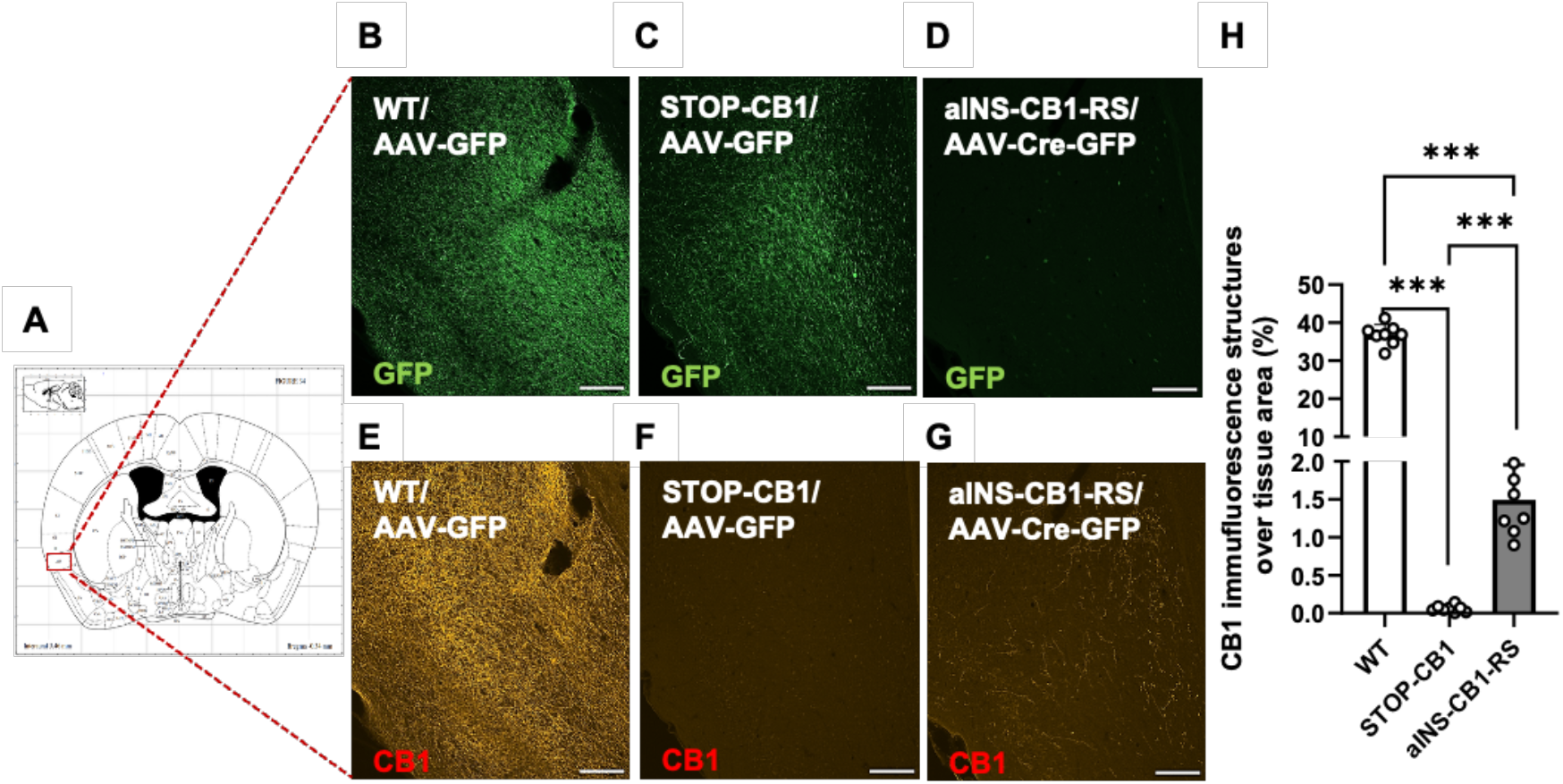
Region-specific rescue of CB1R expression in the aINS. **(A)** Representative coronal section illustrating the posterior insular cortex (pINS) according to the Paxinos and Franklin mouse brain atlas. **(B–D)** GFP fluorescence in the pINS of WT, aINS-CB1-RS, and STOP-CB1 mice following viral injection. **(E–G)** CB1R immunostaining in the same region shows endogenous expression in WT mice, restored CB1R expression in aINS-CB1-Rescue mice, and the absence of signal in STOP-CB1-KO mice. All images represented as maximum intensity projection of 10 μm thick z-stack **(H)** Quantification of CB1R immunofluorescence intensity in the pINS across experimental groups. Data are presented as mean ± SEM (n = 3 WT, n = 2 STOP-CB1, n = 4 aINS-CB1-RS). Statistical analysis was performed using one-way ANOVA followed by Bonferroni post hoc tests. ***p < 0.001. Scale bars: 20 μm.

Given the dense projections from the aINS to the posterior insular cortex (pINS), where presynaptic CB1Rs are expected to localize, we evaluated CB1R immunoreactivity in this projection field. WT mice displayed robust CB1R expression in the pINS, whereas STOP-CB1 mice showed the absence of CB1R signal. Importantly, aINS-CB1-RS mice exhibited restored CB1R immunoreactivity in the pINS, consistent with successful re-expression of CB1R in aINSr projection neurons (Fig. 2B–G). Quantification of CB1 immunofluorescence confirmed a significant genotype effect (one-way ANOVA: F(2, 23) = 1366, p < 0.001), with CB1-KO mice showing markedly reduced signal compared to WT (p < 0.001), and aINS-CB1-RS mice displaying significantly increased CB1R levels compared to STOP-CB1 animals (p < 0.001), approaching WT levels (Fig. 2H). These results demonstrate successful and region-specific restoration of CB1R expression in the aINS and its projection targets, validating the rescue approach.

### 3.2. Restoration of CB1Rs in aINS improves core ASD-like behaviors: reduced repetitive actions and rescued social novelty

Repetitive-like behavior was evaluated using the marble burying test. One-way ANOVA revealed a significant genotype effect (F(2, 93) = 6.16, p < 0.01). STOP-CB1 mice did not differ from WT controls in the number of marbles buried (p = 0.4155). In contrast, aINS-CB1-RS mice buried significantly fewer marbles than both WT (p = 0.0008) and CB1-KO mice (p = 0.017) (Fig. 3A). These results indicate that selective restoration of CB1R in the aINS decreases repetitive-like digging behavior, despite the absence of a baseline difference between WT and STOP-CB1 animals.

**Figure 3.**
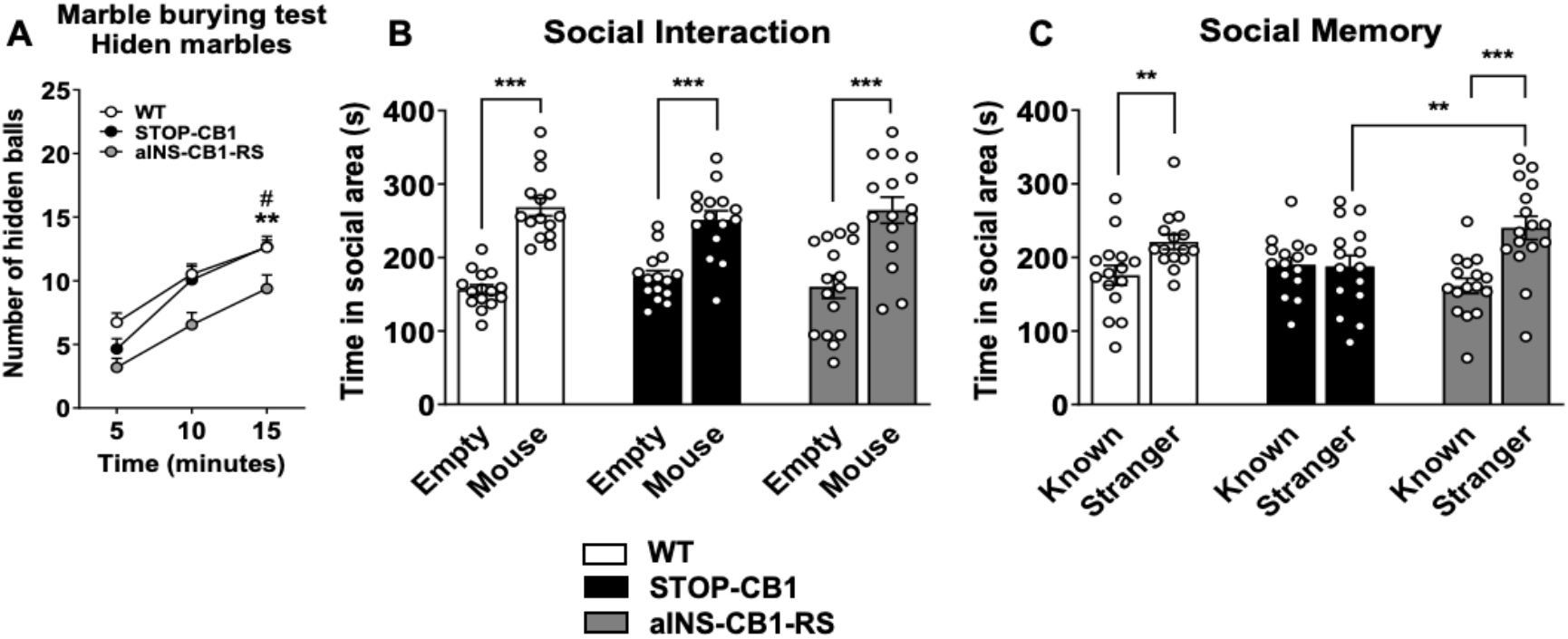
Restoration of CB1 receptors in the anterior insula reduces repetitive-like behavior and rescues social novelty recognition. **(A)** Marble burying test. aINS-CB1-RS mice buried significantly fewer marbles compared to WT (p = 0.0008) and STOP-CB1 mice (p = 0.017), whereas no difference was observed between WT and STOP-CB1 animals (p = 0.4155). Data are presented as mean ± SEM (n = 43 WT, n = 28 CB1-KO, n = 26 aINS-CB1-RS). **(B)** Time spent exploring the unfamiliar mouse versus an empty cage. All groups showed a significant preference for the unfamiliar mouse (p < 0.001), with no genotype differences. **(C)** Social novelty phase for social memory assessment. Following the introduction of a novel conspecific (Stranger), WT (p = 0.04) and aINS-CB1-RS mice (p < 0.001) showed significant preference for the novel over the familiar mouse, whereas STOP-CB1 mice did not (p = 0.9119). Data are presented as mean ± SEM (n = 15 WT, n = 15 STOP-CB1, n = 16 aINS-CB1-RS). Statistical significance was defined as **p < 0.01, ***p < 0.001.

During the sociability phase of the social interaction three-chamber test, all groups showed a strong preference for the chamber containing a conspecific over the empty cage (main effect of chamber: F(1, 43) = 52.47, p < 0.001), with no significant genotype effect or genotype × chamber interaction (F(2,43)=0.597, p > 0.05, Fig. 3B). These findings indicate that basal social approach behavior is not altered by CB1R deletion or insular rescue.

In the social memory phase, a significant genotype × stimulus interaction was observed (F(2, 43) = 3.746, p < 0.05), without a significant effect of genotype (F(2, 43) = 0.802, p > 0.05) and global effect of stimulus (F(1, 43) = 11.033, p < 0.01). STOP-CB1 mice failed to discriminate between the novel and familiar conspecific (paired comparison: p = 0.9119), indicating impaired social recognition memory. In contrast, WT and aINS-CB1-RS mice displayed a significant preference for the novel conspecific (p = 0.04 and p < 0.001, respectively) but not STOP-CB1 mice (Fig. 3C). These data demonstrate that CB1R expression exclusively in the aINS is sufficient to restore social novelty recognition.

### 3.3. Selective CB1Rs re-expression in the aINS reduces anxiety-like behavior without affecting locomotor activity, but fails to prevent depressive-like phenotype

Analysis of spontaneous locomotor activity revealed no significant genotype effect on total horizontal activity (two-way repeated-measures ANOVA: F(2, 87) = 2.939, p > 0.05) (Fig. 4A). Thus, behavioral differences observed in marble burying, social novelty, and anxiety paradigms cannot be attributed to gross motor impairments. Vertical activity (rearing) differed across groups (F(2, 87) = 17.85, p < 0.001), with STOP-CB1 and aINS-CB1-RS mice displaying reduced rearing relative to WT animals (p < 0.001, Fig. 4B).

**Figure 4.**
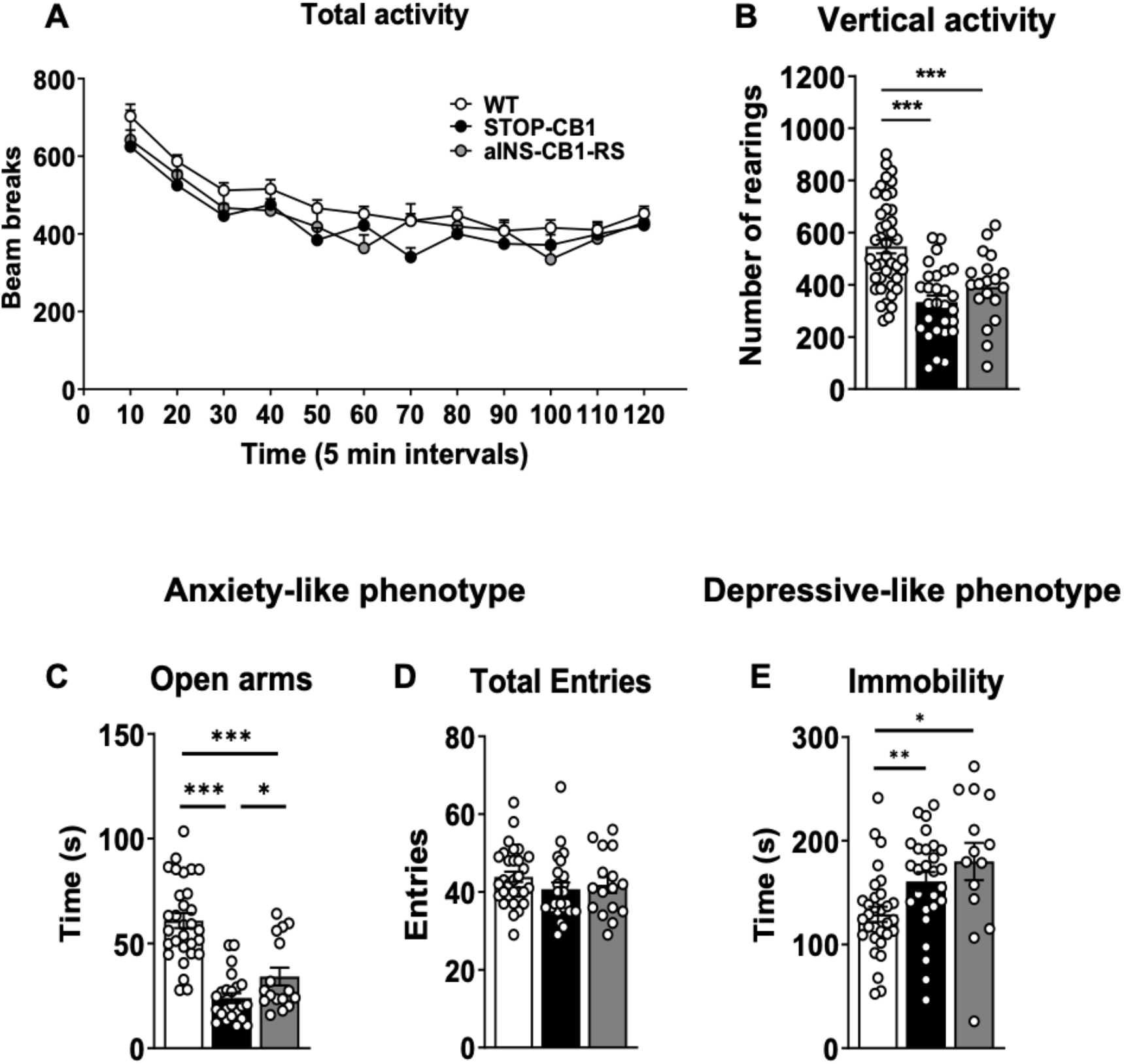
Selective restoration of CB1R in the aINS reduces anxiety-like behavior without affecting locomotor activity, but does not rescue depressive-like behavior. **(A)** Total horizontal locomotor activity measured in automated chambers over 2 h (10-min bins). No significant genotype effect was observed (repeated-measures ANOVA, n.s.). **(B)** Vertical activity (rearing) was measured over 2 h. A significant genotype effect was detected (p < 0.001), with STOP-CB1 and aINS-CB1-RS mice showing reduced rearing compared to WT controls. **(C)** Elevated plus maze (EPM): time spent in open arms. aINS-CB1-RS mice spent significantly more time in the open arms compared to STOP-CB1 mice (p < 0.05), although values remained lower than WT controls. **(D)** Total arm entries in the EPM. No significant differences were observed between groups, indicating comparable locomotor activity during the task. **(E)** Forced swim test: immobility time. Both STOP-CB1 and aINS-CB1-RS mice exhibited increased immobility compared to WT animals (genotype effect, p < 0.01). Data are presented as mean ± SEM (n = 42 WT, n = 28 STOP-CB1, n = 20 aINS-CB1-RS). Statistical significance was defined as *p < 0.05, **p < 0.01, ***p < 0.001.

In the elevated plus maze, data showed a non-Gaussian distribution, and a non-parametric Mann-Whitney U test indicated that STOP-CB1 mice spent significantly less time in open arms than WT controls (p < 0.001), consistent with increased anxiety-like behavior. aINS-CB1-RS mice showed significantly increased open-arm time relative to STOP-CB1 mice (p < 0.05), although values remained lower than WT mice (p < 0.001), indicating only partial rescue after re-expression of CB1R in the aINS (Fig. 4C). Total arm entries did not differ between groups (F(2, 66) = 1.081, p > 0.05), confirming that anxiety-related differences were not secondary to altered locomotion (Fig. 4D).

Depressive-like behavior was not rescued by insular CB1R restoration. In the forced swim test, genotype significantly affected immobility time (F(2, 69) = 6.009, p < 0.01). Both STOP-CB1 (p < 0.05) and aINS-CB1-RS mice (p < 0.01) displayed increased immobility compared to WT controls (Fig. 4E). No significant difference was observed between STOP-CB1-KO and aINS-CB1-RS mice, indicating depressive-like behaviors are not normalized by rescue of insular CB1Rs.

## Discussion

In the present study, we demonstrate that selective restoration of CB1R signaling in the aINS is sufficient to modulate several behavioral domains relevant to NDDs. Region-specific re-expression of CB1R in a CB1R-deficient background (STOP-CB1) was sufficient to reduce repetitive-like behavior, restore social novelty discrimination, and partially normalize anxiety-like behavior while leaving basal sociability and locomotor activity unaffected. In contrast, depressive-like behavior was not rescued by insular CB1R restoration. These findings identify the aINS as a critical CB1-dependent modulatory hub regulating socio-emotional and repetitive behavioral phenotypes.

The insular cortex is increasingly recognized as an integrative center linking interoceptive processing with social and affective behavior^13,15,17^. Functional and structural alterations of the insula have been reported across multiple NDDs^29,30^. Our results extend this framework by demonstrating that local endocannabinoid signaling within the aINS is not merely correlated with these behaviors but is sufficient to influence specific behavioral outputs. Notably, restoration of CB1R expression selectively rescued social novelty recognition, without altering baseline sociability, suggesting that insular CB1R signaling plays a particularly important role in higher-order social memory processes rather than in primary social approach. This aligns with recent evidence that the aINS encodes social novelty and social preference^14,18,19^.

Similarly, the reduction in marble burying behavior following insular CB1R restoration supports a role for the aINS in the modulation of repetitive-like actions. Although marble burying does not exclusively model repetitive behavior, it reflects innate repetitive digging behavior that is frequently altered in rodent models of NDDs^23^. The dissociation between repetitive behavior rescue and the absence of baseline differences between WT and STOP-CB1 animals suggests that region-specific CB1R signaling can actively modulate behavioral output beyond restoring a simple loss-of-function phenotype. The ability of region-specific CB1R restoration to alter this behavior is consistent with prior evidence implicating endocannabinoid signaling in repetitive and compulsive phenotypes^8,31^.

The partial normalization of anxiety-like behavior further indicates that CB1R signaling in the aINS contributes to affective regulation^12,22^. However, the inability of insular CB1R restoration to rescue depressive-like behavior implies that this phenotype likely involves distributed CB1R-dependent circuits beyond the aINS. This dissociation highlights the anatomical specificity of CB1R-mediated behavioral regulation and supports the concept that distinct emotional domains are governed by different segregated neural networks. Specifically, the failure to rescue depressive-like behavior suggests that CB1R-dependent regulation of despair-like responses could involve broader limbic circuits beyond the aINS, such as the prefrontal cortex, amygdala, or hippocampus^12,32^.

Importantly, locomotor activity was largely unaffected by insular CB1R restoration, indicating that behavioral effects observed in repetitive, social and anxiety paradigms are unlikely to reflect generalized motor confounds. These findings demonstrate that CB1R signaling within the aINS is sufficient to regulate selective socio-emotional and repetitive behavioral domains relevant to NDDs

In conclusion, we have therefore identified the aINS as a key region in which endocannabinoid signaling modulates core behavioral dimensions associated with NDD pathology. These findings provide mechanistic insight into region-specific CB1R function and suggest that targeted modulation of insular endocannabinoid signaling may represent a promising avenue for future therapeutic exploration.

## Acknowledgements

This work was supported by CRC1193 “Neurobiology of resilience” to B.L.; ERA-NET Neuron/Bundesministerium für Bildung und Forschung (BMBF)/DLR Projektträger ref. 01EW2203A to B.L. and ref. 01EW2203B to M.J.S. Spanish “Ministerio de Ciencia, Innovación y Universidades, Agencia Estatal de Investigación (AEI)” (PID2020-120029GB-I00/MICIN/AEI/10.13039/501100011033, RD21/0009/0019 and PDI2023-1511680B-C21), the Spanish “Instituto de Salud Carlos III, RETICS-RTA” (#RD12/0028/0023), the “Generalitat de Catalunya, AGAUR” (#2020 SGR), “ICREA-Acadèmia” (#2025) and the Spanish “Ministerio de Sanidad, Servicios Sociales e Igualdad”, “Plan Nacional Sobre Drogas of the Spanish Ministry of Health” (#PNSD-2022) to R.M., “la Caixa Health” LCR/PR/HR22/5240017 to R.M. and E.M-G., “Plan Nacional Sobre Drogas of the Spanish Ministry of Health” (#PNSD-2019I006, #PNSD-2023I040) and Spanish “Ministerio de Ciencia e Innovación” (ERA-NET) PCI2021-122073-2A to E.M-G.

We are very grateful to R. Martín and F. Porrón for their technical support. The authors gratefully acknowledge the IMB Microscopy and Histology CF for their support and assistance in this work.

Figure 1 was created with BioRender.

## Author contributions

E. M.-G. and R. M. conceived and designed the behavioral studies with input from B. L. and I. R. A.; B. L. generated the conditional transgenic mice; A. B. and A. G-B. performed the behavioral experiments and the statistical analyses and graphs with the supervision of E.M.-G. and R.M.; I. R. A performed the immunofluorescence studies. P.T. and I.R.A. performed the confocal microscopy analysis and P.T. performed the subsequent imaging analysis. E.M.-G., P. M.-A., G. H. and R.M. wrote the manuscript and I.R.A and B. L. provided critical review of the manuscript with inputs from all the other authors.

## Data availability

Individual data points are graphed in the main figures. All the relevant data that support this study are available from the corresponding author to any interested researcher upon reasonable request.

## Conflict of interest

The authors declare no conflict of interest.

## Notes

### Competing Interest Statement

The authors have declared no competing interest.

